# Structure and evolution-guided design of minimal RNA-guided nucleases

**DOI:** 10.64898/2025.12.08.692503

**Authors:** Petr Skopintsev, Isabel Esain-Garcia, Evan C. DeTurk, Peter H. Yoon, Zehan Zhou, Trevor Weiss, Maris Kamalu, Ajit Chamraj, Kenneth J. Loi, Conner J. Langeberg, Ron Boger, Hunter Nisonoff, Hannah M. Karp, LinXing Chen, Honglue Shi, Kamakshi Vohra, Jillian F. Banfield, Jamie H.D. Cate, Steven E. Jacobsen, Jennifer A. Doudna

## Abstract

The design of RNA-guided nucleases with properties not limited by evolution can expand programmable genome editing capabilities. However, generating diverse multi-domain proteins with robust enzymatic properties remains challenging. Here we use an artificial intelligence-driven strategy that couples structure-guided inverse protein folding with evolution-informed residue constraints to generate active, divergent variants of TnpB, a minimal CRISPR-Cas12-like nuclease. High-throughput functional screening of AI-generated variants yielded editors that retained or exceeded wild-type activity in bacterial, plant and human cells. Cryo-EM-based structure determination of the most divergent active variant revealed new stabilizing contacts in the RNA/DNA interfaces across conformational states, demonstrating the design potential of this approach. Together these results establish a strategy for creating non-natural RNA-guided nucleases and conformationally active nucleic acid binders, enlarging the designable protein space.

**One-sentence abstract:** An evolution- and structure-conditioned model enables design of active RNA-guided nucleases with new nucleic acid contacts resolved by cryo-EM.

## Introduction

CRISPR-Cas systems have revolutionized genome editing by enabling programmable, sequence-specific DNA and RNA targeting, forming a foundation for precise genetic and epigenetic perturbations (*1*–*7*). Protein design has the potential to extend these capabilities by creating RNA-guided nucleases with functions and properties, including sequence and structures, that are not observed in nature.

However, design of this type is particularly challenging because these nucleases are multi-domain nucleic acid binders whose activity depends on coordinated RNA and DNA recognition, activation and cleavage by distinct conformational states (*8–11*). Sequence-based biological language models (LMs) trained on evolutionary data can generate RNA-guided nucleases by inferring sequence–function relationships, but they often generate nuclease sequences highly similar to the reference sequences they are trained on, even after extensive post-hoc filtering (*12, 13*). Structure-guided rational design approaches offer a robust strategy to sample highly divergent protein sequences, and also structures not found in nature, extending into *de novo* design (*14*–*19*). While this approach has been successful for the generation of dynamic switches and DNA binders (*20*–*22*), the design of complex enzymes such as RNA-guided nucleases bearing multiple functional domains and conformational states, remains an attractive but unresolved challenge.

We reasoned that inverse protein folding models (*17*–*19*), a structure-conditioned approach, coupled with evolution-ary information could generate highly diverse proteins while retaining function. To test the potential of such a design strategy, we selected TnpB, a diverse family of small transposon-encoded ancestors of CRISPR-Cas12 nucleases that mediate RNA-guided DNA cleavage, transcriptional regulation and genome targeting across all kingdoms of life (*23*– *29*). These features render TnpBs as a promising scaffold for exploring new RNA-guided nucleases not produced by natural evolution.

Here we show that structure- and evolution-based conditioning can generate miniature, active RNA-guided nucleases with a high experimental success rate (24%) and without post-hoc filtering. Inverse folding coupled with evolutionary residue constraints predicted highly divergent RNA-guided enzymes, many of which are active in bacterial, plant and human cells. Moreover, cryo-EM based structural analysis of the most diverse yet active variant revealed that this approach can introduce new structural nucleic acid contacts consistent with the protein conformational transitions essential for biochemical function. Together, this work expands the accessible evolutionary space of RNA-guided nucleases and provides a new strategy for the design of non-natural multi-state enzymes.

## Results

### Evolution- and structure-conditioned design of TnpB

We began our study by evaluating in silico whether the ESM Inverse Folding (ESM-IF1) model (18) could redesign the minimal RNA-guided nuclease ISDra2 TnpB to create functional enzymes with non-natural sequences. ESM-IF1-generated sequences accurately recapitulated the input fold predicted by AlphaFold2 (30) (Fig. 1A; fig. S1A), preserved the DED catalytic triad in the RuvC domain (Fig. 1B; fig. S1B), and captured amino-acid variation found among natural homologs (figs. S1C–D) (18). In contrast, LigandMPNN (19, 31) reproduced the TnpB fold but failed to maintain the catalytic triad across all model generation temperatures.

**Fig. 1.**
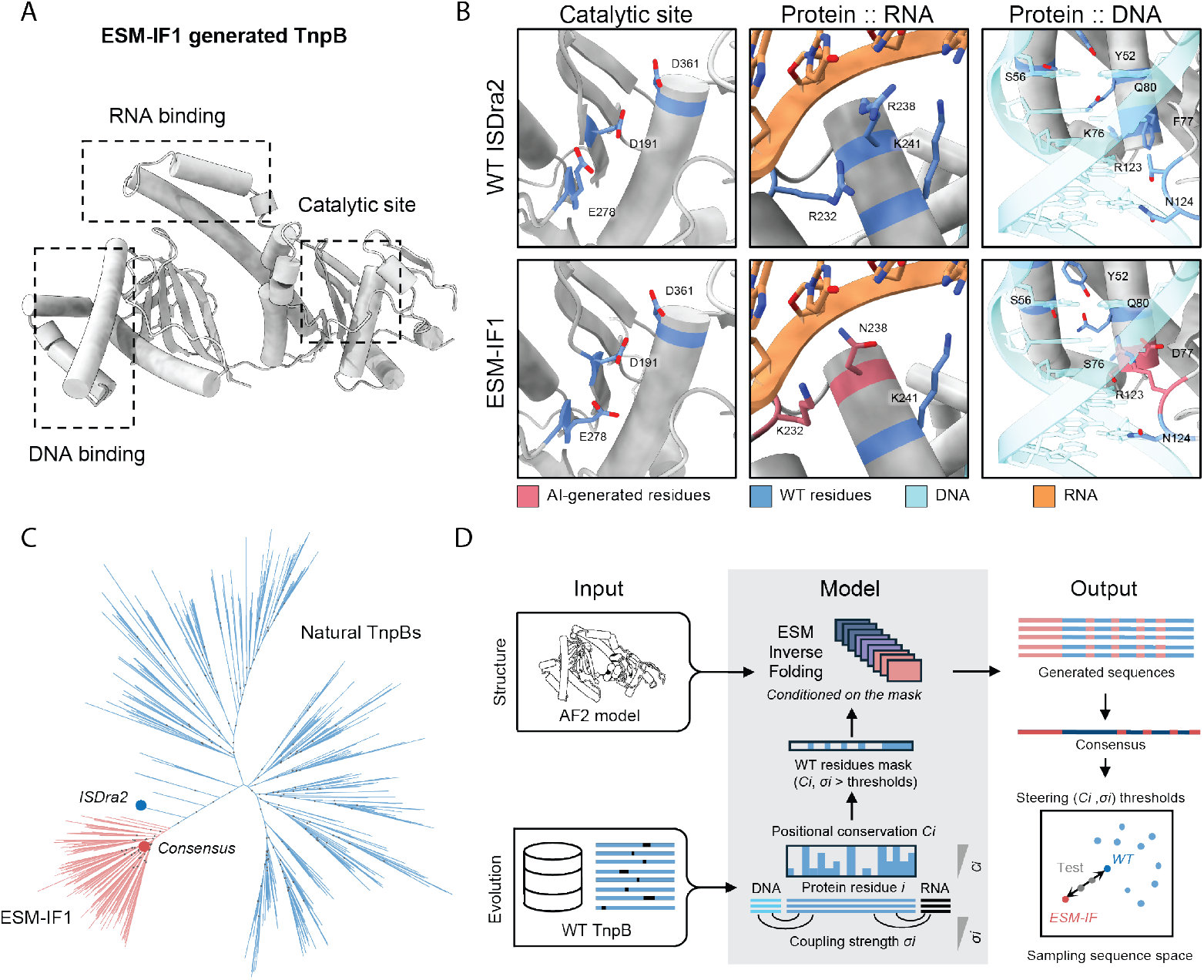
RNA-guided nuclease TnpB design strategy with an ESM Inverse Folding (ESM-IF1) model. **(A)** ESM-IF1 generates sequences that fold into ISDra2 TnpB structure **(B)** RuvC catalytic residues returned by ESM-IF are similar to the WT, whereas those that bind to the nucleic acids vary between WT and ESM-IF sequences. **(C)** Phylogenetic analysis shows that sequences generated by ESM-IF1 cluster into a distinct clade, reflecting substantial divergence from natural TnpBs. Nodes with ≥90% bootstrap support are marked with gray dots. **(D)** In order to preserve ISDra2 TnpB-specific nucleic acid-binding residues in the generated sequences, the ESM-IF model is conditioned on wild-type residues conserved above chosen threshold for positional conservation *C*_*i*_ or co-evolutionary coupling strength *σ*_*i*_ to the ligand nucleic acids. By increasing the (*C*_*i*_, *σ*_*i*_) thresholds, the divergence from the WT ISDra2 TnpB is gradually increased, ultimately yielding fully unconstrained sequences. Consensus sequences at selected parameters are then calculated and tested for activity.

Phylogenetic analysis placed ESM-IF1 sequences in a distinct clade separate from natural TnpBs (Fig. 1C), sharing only 50–60% identity with ISDra2 TnpB (figs. S1E, F). The model produced chemically conservative substitutions across the DNA–RNA interface (Fig. 1B). However, it also introduced non-synonymous mutations at residues K76, F77 and T123, known to mediate nucleobase-specific recognition of the transposon-associated motif (TAM) sequence (32), a short DNA motif priming TnpB activation similar to the protospacer adjacent motif (PAM) of CRISPR-Cas9 (1). Similarly, residue substitutions diverging from the wild-type (WT) ISDra2 TnpB occurred at the interface between the protein and its guide RNA (or reRNA for TnpB). This suggests that ESM-IF1 captures TnpB’s fold and catalytic DED motif and partially recovers nucleic acid-binding residues, but misassigns some contact positions in the absence of a cognate RNA/DNA context.

We therefore explored targeted conditioning to enforce functional protein-RNA and protein-DNA contacts. Typically, inverse-folding models fix residues within a defined radius of experimentally resolved bound ligands (33). For TnpB, however, the 49 kDa RNA–DNA heteroduplex binds the 46 kDa protein over a large interface, requiring many residues to be fixed, and leaving little space to redesign. Furthermore, for dynamic proteins such as RNA-guided nucleases, available cryo-EM structures (32, 34) may miss transient protein-DNA/RNA contacts, which may result in not fixing all residues necessary for function.

To address this, we developed a masking strategy for inverse folding that introduced functionally critical residues from evolutionary data. Previous bioinformatic studies showed that DNA- and RNA-binding residues can be identified from phylogenetic conservation or co-evolution signals (35, 36). We thus derived positional conservation *C*_*i*_ values for TnpB residues *i* from multiple-sequence alignments of natural TnpBs, and coupling signals *σ*_*i*_ from a Potts model (GREMLIN) (37) trained on either paired TnpB–RNA or TnpB-DNA sequences mined from genomic databases (Methods; fig. S2). Residues exceeding given *C*_*i*_, *σ*_*i*_ thresholds *C*_*0*_, *σ*_*0*_ were fixed to their wild-type identities in ISDra2 TnpB, producing sequence masks for conditional ESM-IF1 generation. Varying thresholds *C*_*0*_, *σ*_*0*_ effectively tuned the balance between structural and evolutionary input to the model (Fig. 1D). We further found that the consensus averaging of generated sequences had lower model perplexity than individual sequences, indicating improved fold-compatibility and possibly higher chances of activity (fig. S3).

### High-throughput bacterial screens identify active variants

Our design objective was to experimentally determine which (*C*_*0*_, *σ*_*0*_) combinations produced active nucleases while minimizing fixed residues, thereby maximizing the sampled sequence space and exploring synthetic TnpB diversity. To maximize the generation rate of active proteins, we tested consensus sequences derived from 10,000 ESM-IF1 generations per *C*_*0*_ or dual (*C*_*0*_, *σ*_*0*_) conditions. To screen the generated TnpB variants for activity, we employed a bacterial assay in which cell recovery is proportional to TnpB-mediated cleavage of a plasmid carrying the ccdB toxin gene (38) (Fig. 2A-C).

**Fig. 2.**
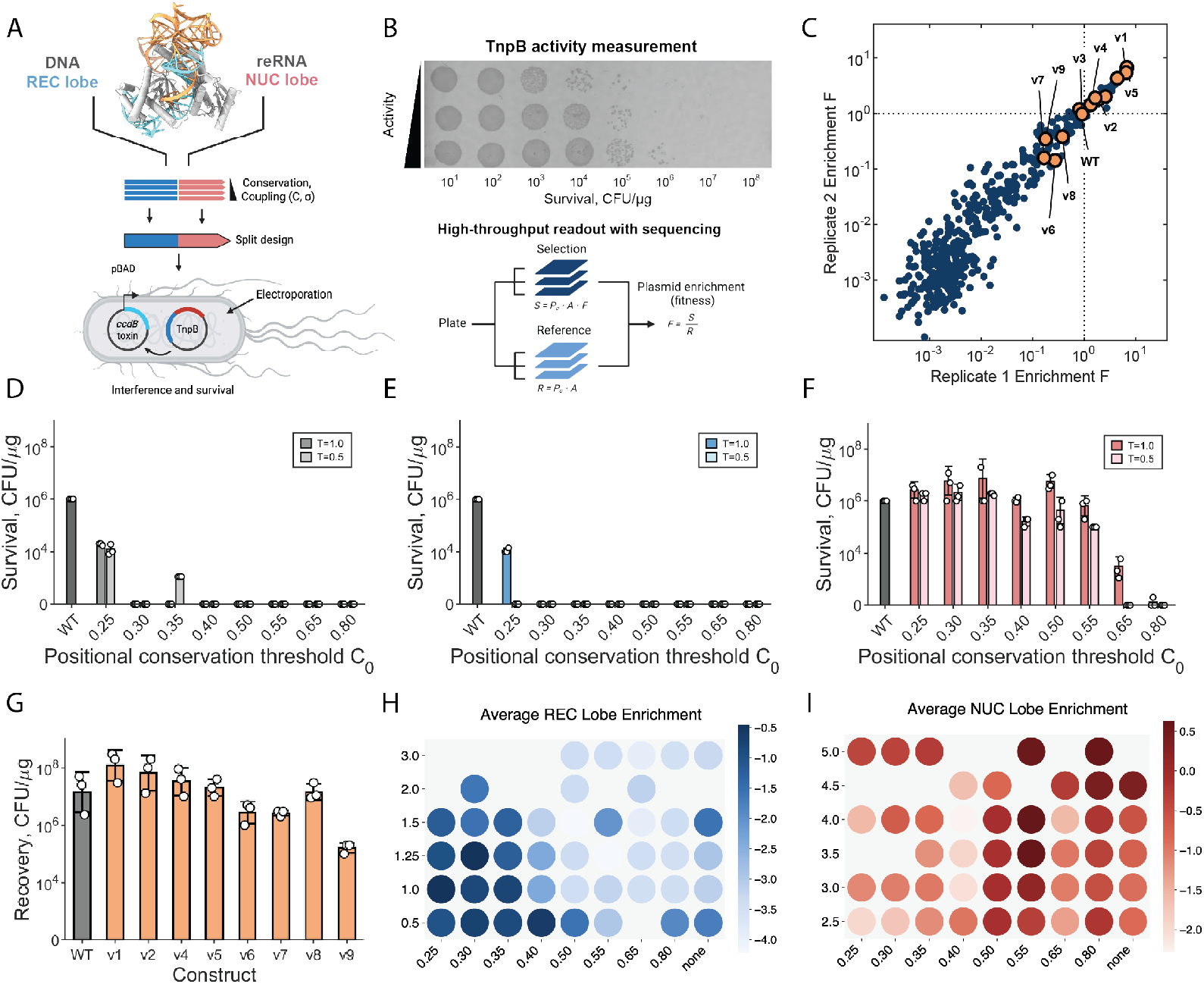
Screening AI-generated TnpB variants conditioned on positional conservation (*C*_*i*_) and coupling strength (*σ*_*i*_) thresholds. **(A)** Splitlobe design of TnpB allowed combinatorial testing of REC and NUC lobes. Plasmids expressing TnpB variants were electroporated into *E. coli* cells carrying an arabinose-inducible toxin (pBAD-ccdB). Active variants cleave the toxin plasmid, enabling survival on selective agar. **(B)** Activity was measured by spot plating (top) or by pooled bacterial selection coupled to high-throughput sequencing (bottom). **(C)** Log_10_ enrichment of generated variants across two experimental replicates. Data were normalized such that reference WT ISDra2 TnpB enrichment = 1. **(D)** Activities of full-length variants generated across *C*_*i*_ thresholds at two ESM-IF1 sampling temperatures. Bars show mean ± SD from three replicates. **(E, F)** Cleavage activities of REC (E) and NUC (F) lobes tested individually with their WT counterparts. **(G)** Recovery of top variants (1,980 tested in (C)) assembled from REC and NUC lobes conditioned on both (*C*_*i*_, *σ*_*i*_) thresholds. **(H, I)** Heat maps showing enrichment of AI-generated REC (H) and NUC (I) lobes over (*C*_*i*_, *σ*_*i*_) thresholds, averaged over all fused combinations.

Using spot-plating to test full-length proteins generated for a range of conservation (*C*_*i*_) thresholds (*C*_*0*_) and model temperatures (*T*), we observed that three variants at *C*_*0*_ =0.25 and 0.35 exhibited activity, with survival levels of 10^3^-10^4^ CFU/µg, albeit lower than the 10^6^ CFU/µg for the WT (*C*_*0*_= 0.0; Fig. 2D). We hypothesized that as TnpBs are multi-domain proteins, the different lobes may not tolerate comparable mutational depth or achieve similar generation success. Therefore, we implemented a split-domain strategy in which the generated DNA- and RNA-binding domains, referred to as REC and NUC respectively, were experimentally tested separately.

Inspired by the domain-swapping strategy previously applied to CRISPR–Cas9 engineering (39), we fused the AI-generated lobes to their WT counterparts to assess them individually (Fig. 2E, F). Supporting our hypothesis, the lobes exhibited different tolerances to sequence divergence. Only one AI-generated REC lobe variant at *C*_*0*_ = 0.25 was functional when fused to the WT NUC lobe, whereas 13 out of 16 NUC-lobe variants retained activity, often higher than the WT for *C*_*0*_ thresholds as high as 0.65. These results suggest that the NUC lobe accepts higher levels of divergence from the WT, whereas the REC lobe is more sensitive to substitutions under single-parameter (*C*_*i*_) conditioning.

To sample a broader range of substitutions and further increase the diversity of active variants, we tested a dual-conditioning strategy by fixing residues according to both positional conservation (*C*_*i*_) and protein–nucleobase coupling strength (*σ*_*i*_). REC and NUC lobes were independently generated under (*C*_*O*_, *σ*_*O*_) thresholds (Methods; figs. S2, S4-5) and assayed in all pairwise REC–NUC combinations (44 × 45 lobes; 1,980 fusions) using a high-throughput pooled library selection assay (Fig. 2B). Activity was quantified as the enrichment of lobe-pair plasmids under selective (+arabinose) versus non-selective (–arabinose) conditions.

In contrast to single-parameter (*C*_*i*_) conditioning, dual (*C*_*i*_, *σ*_*i*_) masking produced a markedly broader distribution of active variants, revealing more diverse and functional REC and NUC lobes (Fig. 2H, I). Of the 466 active lobe combinations (24% of 1,980 designs), approximately 8% exceeded the activity of the WT (Fig. 2C). The most diverse and active variants identified in the pooled assay (v1-v9) were validated by spot-plating (Fig. 2G) and were shortlisted for testing in human and plant genome editing assays.

### AI-designed TnpB enzymes are active genome editors in human and plant cells

We next evaluated the genome editing activity of representative TnpB variants (v1-v9), spanning high activity and sequence diversity ranging from 77-91%, and WT ISDra2 in an endogenous blue fluorescent protein (BFP) gene knockout assay in HEK293T cells using biological triplicates (Fig. 3A; fig. S6). The WT enzyme exhibited an average of 28% editing. Variants v2-4 and v6-8 showed comparable values to the wild-type in the range of 23-32%. Importantly, v1 and v5 showed the highest increases in editing efficiencies over WT, with values of 46% (p-value<0.001) and 50% (p-value<0.001), respectively (Fig. 3B). These results confirmed the successful generation of active and diverse variants.

**Fig. 3.**
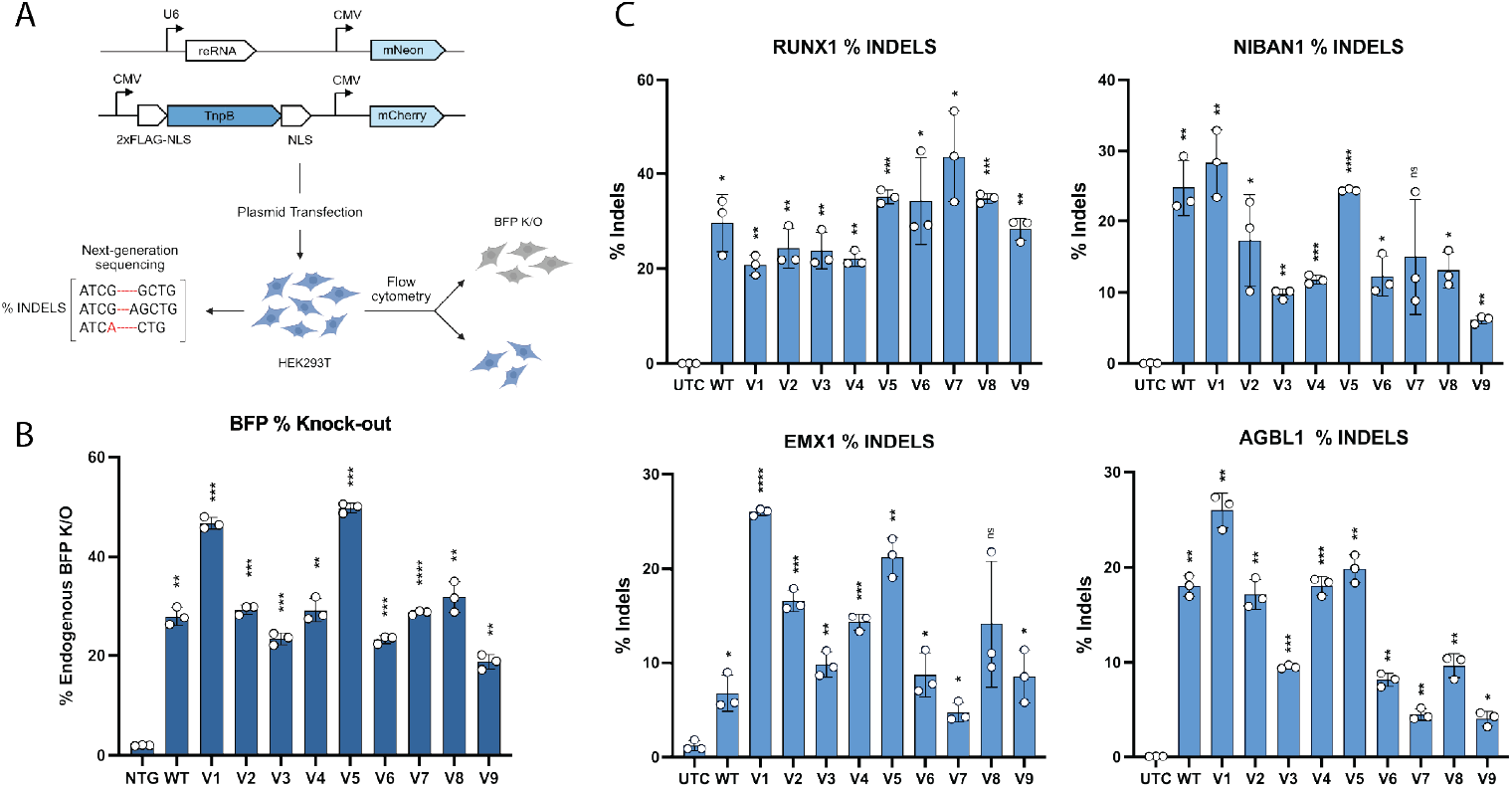
AI-generated TnpB genome editing in HEK293T cells. **(A)** Plasmids encoding variant proteins and corresponding guide reRNAs were co-transfected into HEK293T cells. Editing efficiency was assessed by either **(B)** *BFP* gene knockout measured by flow cytometry or **(C)** next-generation sequencing (NGS) quantification of indels at endogenous loci (*RUNX1, NIBAN1, EMX1, AGBL1*). Bars show mean ± SD from three replicates. P-value: ns > 0.05, * ≤0.05, ** ≤0.01, *** ≤0.001, **** ≤0.0001.

Next, to assess programmability and editing potential, we targeted endogenous human and plant genomic loci. Four endogenous genes interrogated previously, RUNX1, NIBAN1, EMX1, and AGBL1, were targeted by the AI-generated TnpBs in HEK293T cells (23, 40), and the percentage of insertions and deletions (indels) was calculated by next-generation sequencing (NGS). While most variants exhibited similar activity to the WT, v1 and v5 again were the most active variants across the tested loci. Variant v1 showed editing of 26%, 26%, 28% and 21% for EMX1, AGBL1, NIBAN1 and RUNX1, while v5 showed 21%, 20%, 24% and 35%, compared to 7%, 18%, 25% and 30% for the WT (Fig. 3C). Notably, v1 and v5 showed respective 3.8- and 3.1-fold increases over WT at the EMX1 locus (v1 p-value <0.0001, v5 p-value <0.01). Due to their minimal size, TnpBs are well suited for plant genome editing by viral vector delivery (41). Thus, we tested the nine selected variants in Arabidopsis protoplast cells by targeting the AtPDS3 gene at four distinct sites. Consistent with the HEK293T results, v1 outperformed ISDra2 at nearly all tested targets (fig. S7).

### Conformational dynamics provide mechanistic insights into TnpB activity

The highly divergent variant v7 showed the highest absolute editing efficiency in our cellular experiments (44% at RUNX1; Fig. 3C). This variant shared 77% sequence identity with the WT (excluding the unstructured C-terminus dispensable for activity, which was kept fixed; 83% for the REC lobe, 72% for the NUC lobe). In line with our design objective, approximately one in four residues in v7 was generated by the evolution-conditioned ESM-IF1 model, totaling 85 new positions. Many of these residues lined the nucleic acid interface alongside masked WT residues, and included both conservative and non-conservative substitutions (fig. S8). These results prompted us to examine the structural basis of how AI-introduced substitutions facilitate the TnpB cleavage mechanism. To study how the newly generated residues facilitate function in the AI-generated protein v7, we determined its ternary complex structure with cryo-EM, capturing TAM-bound and R-loop-formed states set at 2.8 Å resolution (Fig. 4; fig. S9).

**Fig. 4.**
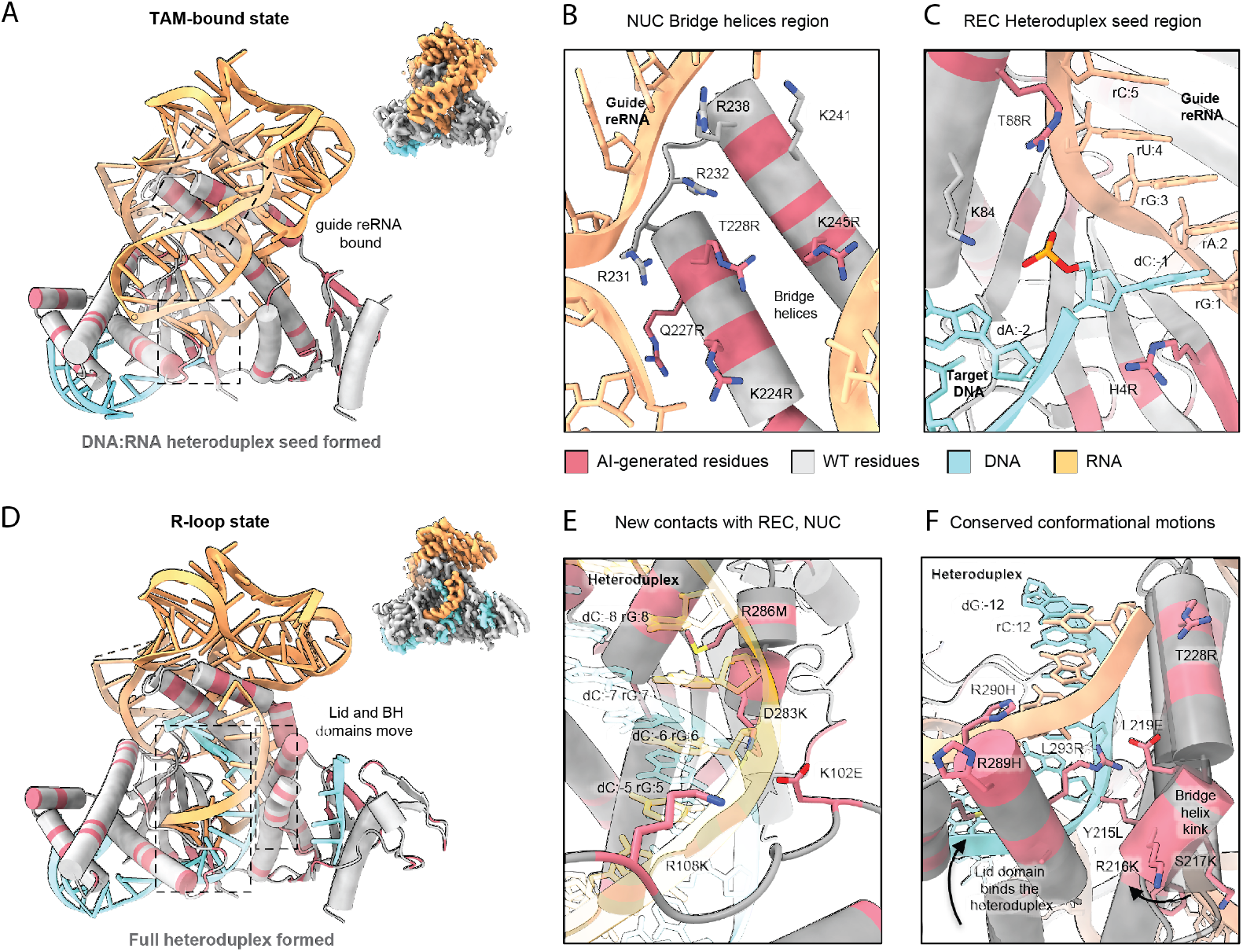
Cryo-EM structures of the AI-generated variant v7 reveal new RNA–DNA interface residues and conserved conformational motions. **(A)** Cryo-EM reconstructions of the TAM-bound (heteroduplex seed formed) and **(D)** R-loop (full heteroduplex) states at 2.8 Å resolution. **(B)** In the NUC lobe, newly introduced residues (coral) form new contacts with the guide reRNA. **(C)** The phosphate lock residue K84 stabilizes the first DNA base pairing with the RNA spacer; new putative contacts H4R and T88R appear near the heteroduplex seed. (**E**) Both REC and NUC lobes exhibit additional AI-generated contacts with the RNA:DNA heteroduplex. **(F)** Conserved conformational motions include lid-domain stabilization and bridge-helix kinking mediated by AI-generated residues (L289R-L293R, and Y214F–L219E). Color scheme: AI-generated residues, red; fixed WT residues, gray; RNA, orange; DNA, cyan.

For both states, in the NUC lobe, the RNA–protein interface exhibited a build-up of positive electrostatic potential facilitated by newly introduced residues K224R, Q227R, T228R and K245R (Fig. 4A, B). The RNA was further stabilized by the masked WT residues R232, R238 and K241, which interact with co-evolving RNA nucleobases. Additional novel contacts at K256Q and R270K lined the RNA pseudoknot region. Notably, our cryo-EM analysis resolved a TAM-bound conformational intermediate state not previously reported for TnpBs. In this new state, the protein and RNA conformations closely resembled the binary complex of ISDra2 TnpB (8BF8 in (34)), but the complex was also bound to DNA, containing a single nucleotide pair formed in the guide reRNA–target DNA heteroduplex seed region. While the majority of the WT protein-DNA contacts were kept due to strong co-evolution with the TAM motif, novel N4R residues stabilized the heteroduplex seed pair together with the conserved “phosphate lock” K84 (42) (Fig. 4C). Unlike in the ISDra2 structure (34), reRNA Stem 1 was structured and bound to the REC lobe, likely stabilized by the new I40K salt bridge with the phosphate backbone (fig. S10).

Our R-loop-formed state closely resembled the previously reported ISDra2 TnpB ternary complex (8EXA in (34)), with the heteroduplex fully formed along the RNA spacer nucleotides 1-12 (Fig. 4D). The REC lobe positions the AI-generated T88R, K102E, R108K, R113K and R160K residues in contact with the heteroduplex. Further, the lid domain which caps the substrate binding cleft was structured on the heteroduplex through non-natural D283K, R286M, R289H and L293R contacts (Fig. 4E). Remarkably, a helical segment largely composed of ESM-IF1-generated residues Y214F, L215Y, R216K, R217K and L219E preserved the previously observed kink motion that facilitates heteroduplex formation (Fig. 4F) (34). Finally, the NUC lobe was fully structured and bound the ssDNA substrate in the RuvC catalytic pocket, which remained lined by residues found in the WT enzyme.

Together, these structures reveal that the AI-generated residues introduced new electrostatic and hydrogen-bonding networks likely to stabilize the RNA/DNA interface in different protein conformational states. The largely AI-generated segment of the protein preserved the native helix kink, consistent with the protein conformational transitions. Finally, our design approach enabled us to capture a previously unobserved TAM-bound state which may be transient in the WT but is populated in the AI-generated variant.

## Discussion

TnpB-family enzymes, which are compact RNA-guided nucleases ancestral to CRISPR-Cas12, are an attractive target for protein design because they couple programmable DNA targeting and a variety of natural functions to a minimal architecture. However, designing such systems has remained limited because sequence-based biological language models produce proteins that remain close to natural sequences, whereas rational design requires explicit programming of the conformational mechanism.

Here we show that combining a structure-guided inverse folding model with fixing residues derived from TnpB evolutionary data yields miniature, highly divergent TnpB nucleases that retain or exceed WT activity across human, plant and bacterial systems. Our semi-rational design strategy generated 466/1,980 designs (24%) with detectable activity, 8% of which exceeded WT activity. This was achieved by fixing a set of residues identified by evolutionary conservation and protein–nucleobase coupling to retain positions critical for function, allowing us to sample protein sequences divergent from natural homologs.

Unlike LMs that generated proteins that retained WT DNA-binding domains with >99% identity to natural homologs (fig. S11) (12, 13), we created DNA- and RNA-interacting lobes with new contacts that had 83% and 72% identity to their closest counterparts in nature, respectively. The split-design approach further revealed asymmetric tolerance to sequence change: the DNA-recognition REC lobe was markedly more substitution-sensitive than the RNA-binding catalytic NUC lobe. We attribute this sensitivity to finely tuned biophysical requirements and stricter evolutionary constraints on TAM search and recognition (43), consistent with the need for separate optimization of the REC and NUC lobes. This modular design principle could generalize to other multi-domain proteins. Our masking strategy is agnostic to prior experimental structures and is based on evolutionary information. It preserves critical contacts yet permits emergence of new contacts that create conformation-specific RNA/DNA interactions across enzymatic states, as supported by our first insight into the AI-generated RNA-guided nucleases cryo-EM structures.

Supporting the outcome of our design strategy, recent deep mutational scanning (DMS) and directed-evolution studies identified that our AI-generated N4R and Q227R substitutions increase ISDra2 TnpB activity (44, 45). Further, while recent AI-guided engineering studies have used inverse folding models to enhance natural proteins with sin-gle-point mutations (46, 47), our distinct strategy integrates phylogenetic conservation with protein–RNA/DNA co-evolution, enabling the design of highly divergent non-natural proteins.

The appearance of new DNA/RNA contact residues and their alignment with TnpB’s conformational motions raise the possibility that ESM-IF1 may encode emergent biophysical constraints beyond static sequence-structure mapping. Refined by evolutionary constraints in our approach, this capability may be useful for structure-guided design of other multi-state proteins or nucleic acid binders that are represented in genomic databases together with their cognate DNA/RNA sequences (48, 49). Further, it has been proposed that proteins function and evolve as ensembles of conformations rather than as single static structures (50). In that context, the previously unresolved TAM-bound state we observe likely represents a distinct kinetic intermediate relative to the WT TnpB, and provides a precedent for isolating and studying transient conformations that RNA-guided nucleases may sample in nature and beyond. Establishing the foundation for design of RNA-guided systems and flexible nucleic acid binders with an evolution- and structure-guided model, we envision extending their functions beyond natural evolutionary constraints towards de novo design, expanding the designable protein space.

## Acknowledgements

We thank members of the Doudna laboratory and the Innovative Genomics Institute for helpful discussions; UCSF for giving us access to the high-performance computing cluster Wynton to meet our computational needs. D. Bulkley and G. Gilbert at the UCSF CryoEM Core are acknowledged for the support in ternary complex screening; D. Toso, K. Sharma, J. Remis, P. Tobias at the Cal-Cryo at QB3-Berkeley for data collection support; N. Krishnappa for running the NGS core at the IGI; C. Hsu and D. Savage for feed-back on the initial manuscript. S. Chitrananda for discussions on the Potts model. L.E. Valentin-Alvarado for discussions on phylogenetics; B.W. Thornton for discussions on the library vector design; K. Chen and K. Wasko for providing the BFP HEK293T cell line; K. Zhou and J. Ye for invaluable technical and organizational support of the Doudna laboratory; K. Lucas for outstanding leadership and coordination in managing the laboratory’s operations and research activities. Schematic illustrations in Figures 2A and 3A were created with BioRender.com. This paper was typeset with the bioRxiv word template by @Chrelli: www.github.com/chrelli/bioRxiv-word-template

## Author contributions

Conceptualization and methodology: PS, JAD.

Computational design: PS, HN.

Bioinformatics: PS, IEG, CJL, LXC, PHY.

Cloning and bacterial assays: ECD, PS, IEG, AC, PHY, KL, HS, KV. HEK293T

experiments: IEG, PS, PHY, ECD, HMK.

Data analysis: PS, IEG, ECD, RSB.

Plant editing and analysis: TW, MK, SEJ.

Protein purification: ZZ, PS.

Cryo-electron microscopy: PS, IEG.

Writing: PS, IEG, ECD, JAD with contributions from all authors.

## Data and materials availability

TAM-bound and R-loop formed cryoEM states models are deposited in the PDB under accession numbers 9YYG, 9YYH. The CryoEM maps are available in the Electron Microscopy Data Bank under accession numbers EMD-73644, EMD-73645. Sequences of synthetic oligos generated in this study are provided in data S2. Plant gene editing amp-seq data will be accessible at NCBI Sequence Read Archive.

## Materials and Methods

### ESM Inverse Folding (ESM-IF1) sequence generation

The ESM-IF1 inverse-folding model was obtained from GitHub (https://github.com/facebookresearch/esm/tree/main/examples/in-verse_folding), with scripts adapted from the accompanying Google Colab notebook (https://colab.research.google.com/github/facebookre-search/esm/blob/master/examples/inverse_folding/notebook.ipynb). Because experimental structures of wild-type (WT) ISDra2 TnpB contain unstructured regions (32, 34), an AlphaFold2 (AF2) model of ISDra2 was used as input to ESM-IF1 for sequence generation and analysis. For comparison with LigandMPNN, a chimeric atomic model was constructed by merging the AF2-predicted NUC lobe (residues 180–408) with the REC lobe, and nucleic acids from the experimental structure of PDB 8EXA (30, 34).

For each model and temperature condition, 50,000 sequences were generated. Representative sequences were refolded using AF2 and visualized alongside the original AF2 model (fig. S1A). Multiple sequence alignments (MSAs) were performed with Clustal Omega in Geneious Prime 2023.0.4 (https://www.geneious.com/) to analyze residue distributions (fig. S1B). Custom Python and MATLAB 2023a scripts were used to generate substitution matrices (18) (fig. S1C), cross-matrix correlation plots (fig. S1D), distributions of mutual sequence identities (fig. S1E), and recovery analyses showing average sequence identity to the WT ISDra2 as a function of generation temperature (fig. S1F).

For consensus sequence derivation, five randomly sampled batches of 10,000, 3,000, 2,000, 1,000, and 1 sequences from the 50,000 generated dataset were averaged by selecting the most frequent amino acid at each position, yielding five consensus variants. The identities of these averaged sequences relative to the WT (fig. S2A) and their perplexity scores reported by ESM-IF1 (fig. S2B) were calculated, and mean and standard deviation across batches were plotted. Perplexity values converged across temperatures, paralleling the identities of the full 50k consensus sequences. Two-dimensional multidimensional scaling (MDS) embeddings of 1,000 sequences generated at T = 0.5 were produced using a custom Python script (fig. S3C).

### Designing and implementing the positional conservation metric

Protein sequences were retrieved from InterPro (IPR021027, IPR001959, IPR010095) and RefSeq (NF018933.4, NF023742.4, NF038281.1, NF040570.1) based on their HMM families, along with IS605 and IS607 family proteins from ISfinder (http://www-is.biotoul.fr;(*49*)) as of March 11, 2024. Pairwise alignments with the ISDra2 TnpB sequence were performed, and proteins spanning amino acids 5–377 with ≥40% sequence identity were selected. TnpB sequences were downsampled using MMseqs2 at >70% identity and >80% coverage thresholds (*50, 51*). Cluster representatives were aligned to ISDra2 TnpB through five iterations of Clustal Omega and manually curated to include variants that contained expected catalytic residues D191, E278, and D361 and structurally intact using ESMFold (*52*). Curated clusters were expanded back to the full list using custom Python scripts, downsampled again with MMseqs2 (identity >90%, coverage >80%), and re-aligned over 25 iterations of Clustal Omega. Gaps and insertions relative to ISDra2 were removed, and positional conservation frequencies were calculated as *C*_*i*_ = *max*_*rr,i*_), where _*fr,i*_ represents the frequency of amino acid *r* at position *i*.

Masks were generated by fixing WT ISDra2 TnpB residues at positions with *C*_*i*_ above threshold *C*_*o*_, and introducing gaps elsewhere. *C*_*o*_ values of 0.25, 0.30, 0.35, 0.40, 0.50, 0.55, 0.65, and 0.80 were chosen such that each step corresponded to a roughly 5% change in sequence identity relative to the WT. All masks also fixed WT residues at Y52, S56, K76, F77, Q80, T123, and N124, based on mutagenesis data (*32*). These masks were provided to ESM-IF1 in conditional generation mode together with the ISDra2 AF2 model at generation temperatures *T* = 0.5 and 1.0.

### Plasmid construction

Custom plasmids were constructed through Golden Gate Assembly using purified PCR products or custom-designed Gene Fragments (Twist) containing the TnpB REC and NUC lobes encoded by residues 1-179 and 179-377, respectively. All generated TnpBs were co-expressed alongside their reRNA on a single plasmid containing a p15A origin of replication and a *camR* cassette. To allow expression of complexes in both bacterial and mammalian cells via use of a single vector, dual promoter systems were used, with TnpB protein coding sequences cloned under the control of T7 and CMV promoters and reRNAs cloned under the control of T7 and U6 promoters. All synthetic TnpB sequences were human codon optimized using a custom python script. The truncated Trim2 reRNA (*32*) with a spacer sequence designed to work in both bacterial and mammalian activity experiments was used in all cases.

### Plasmid ccdB activity assay

Targeted nuclease activity of TnpB variants was measured using a bacterial selection assay based on cleavage of the ccdB toxin gene (38). A plasmid carrying ccdB under the arabinose-inducible pBAD promoter and an ampicillin resistance cassette was introduced into electrocompetent E. coli BW25141 (*λ*DE3) cells. Transformants were selected on LB agar containing ampicillin and 0.2% glucose to repress toxin expression, and bulk electrocompetent stocks were prepared from liquid cultures.

For activity assays, 100 ng of each TnpB/reRNA plasmid was electroporated into the ccdB-expressing strain, recovered for 2 h at 37 °C in SOB medium, pelleted at 5,000 rpm for 2 min, resuspended in 50 µL SOB, and serially diluted tenfold. Aliquots (1/10 of each dilution) were plated on LB agar supplemented with chloramphenicol (25 µg/mL) and either 0.2% arabinose or 0.2% glucose. Colony-forming units (CFUs) were counted from the most dilute plate with discrete colonies, multiplied by the dilution factor, and normalized to CFU per microgram of DNA. Reported activities represent means of three biological replicates. Bacterial spot-plating assays are shown in fig. S12.

### Designing and implementing the protein-nucleic acid co-evolution metric

To build an alignment of TnpB proteins with their cognate TAM DNA and reRNA sequences, the Genome Taxonomy Database (GTDB, release 220; (53)) was used. Representative IS605 and IS607 TnpB protein sequences from ISfinder were folded using AlphaFold2, and their MSA was generated from DALI structural alignment outputs refined with T-Coffee (54). The alignment, truncated to residues D191–D361 of ISDra2, was used to construct an HMM profile with the HMMER suite (http://hmmer.org/), which was applied to all GTDB coding sequences translated with esl-translate to identify TnpB homologs and their genomic coordinates.

A covariance model (CM) for the IS605 reRNA motif was built using sequences retrieved from NCBI (accessions CP050120.1, NZ_JAGGKC010000000.1, CP000359.1, CP122363.1, CP063232.1, CP126078.1, and CP015438.1), manually aligned in Stockholm format using the ISDra2 TnpB–reRNA complex as a structural guide. The seed alignment was used to generate an initial CM with Infernal (55), which was iteratively refined against the GTDB bacterial reference genome dataset (E-value < 1E−5) through repeated searches, realignment, and manual curation until convergence (fig. S3A).

TnpB loci identified by HMM search were retrieved with 5-kb flanking regions, and the CM was applied to detect associated reRNA sequences. After curation, TnpB and reRNA sequences were aligned using hmmalign and cmalign, respectively, merged into a unified protein–RNA alignment, and trimmed to remove insertions relative to ISDra2 reference sequences. A total of 16,335 protein–RNA pairs were used to train a Potts model in the GREMLIN Colab notebook with a custom extended protein–RNA alphabet (37), and the resulting average product correction (APC) matrices were extracted and analyzed (fig. S3B).

The right end of each parsed reRNA sequence, marked by the conserved “UCAA” motif, was used to identify the right boundary of the transposable element (TE) associated with each TnpB, which in turn facilitated detection of the corresponding left end and the TnpB-specific TAM DNA motif (*56*). To locate the left ends, 1,000-nt flanking sequences surrounding a 1,550-nt region upstream of the reRNA right end (the approximate TE locus) were merged and queried using blastn against a local GTDB instance (*57*). Hits with E-values <1E−10 corresponding to TE insertion sites were aligned to their respective queries, and transposon termini were identified and manually curated. The five nucleotides immediately adjacent to the TE left end were designated as the TAM sequence and paired with their corresponding TnpB, yielding 6,479 TAM–TnpB pairs. This data was subsequently used to train a second Potts model, from which average product correction (APC) matrices were extracted. In TAM–TnpB and TnpB-reRNA multiple sequence alignments, protein and nucleic acid coordinates were denoted *i* and *j*, respectively. Each APC_ij_ value was normalized by its standard deviation to yield *σ*_*ij*_ *= APC*_*ij*_ */ stdev(APC*_*ij*_), providing a standardized co-evolution coupling measure across residue-nucleotide pairs.

For each protein residue *i*, the maximal normalized coupling value *σ*_*ij*_ across all nucleic acid positions *j* was defined as its “coupling strength” *σ*_*i*_ *= max*_*j*_(*σ*_*ij*_) (fig. S3C). Analogous to masking by conservation thresholds (*C*_*o*_), residues with coupling strength values above a defined threshold (*σ*_*o*_) were fixed as WT in subsequent mask generation. *σ*_*i*_ values were derived independently from the TAM–TnpB and TnpB–reRNA datasets for the REC and NUC lobes, respectively (figs. S3C, D).

In the REC lobe, the strongest co-evolution signals (*σ*_*i*_ > 2.5) were observed at residues Y52, G53, S56, S57, D75, K76, F77, Q80, N81, K84, Q121,T123, N124, N125, and Q147, which notably encompassed TAM-proximal positions (Y52, S56, K76, F77, Q80, T123, N124) previously identified as functionally essential by alanine scanning (*32*) and observed to line the dsDNA major groove in experimental TnpB structures (fig. S3D). In the NUC lobe, the strongest coupling with reRNA (*σ*_*i*_ > 4.5) occurred at K143, G146, and Q147 in the REC lobe (fig. S3F) and at K220, R232, A237, R238, K241, and D259 in the NUC lobe (fig. S3G). Given their strong coupling and spatial proximity to reRNA, residues K143, K145, G146, and Q147 were fixed as WT in all subsequent design iterations.

Cross-lobe coupling between the REC and NUC domains was detected at three residue clusters (fig. S4): I162–H165–E168–T244–A247–K251– N255–V254, V8–Q258, and K5–I177– W313; these positions were also masked in future designs. Limited mutagenesis of the *C*_*i*_ = 0.35 (T = 0.5) REC-lobe variant (Fig. 2D) revealed that reverting ESM-IF1 substitutions K95R, D114T, and K155L improved activity, whereas back-conversion V162I decreased activity; accordingly, these residues were fixed as WT in all subsequent REC-lobe designs (fig. S5). Final ESM-IF1 masks were generated using *σ*_*o*_ thresholds ranging from 0.5 to 3.0 for the REC lobe and 1.5 to 5.0 for the NUC lobe (Figs. 2H, I; fig. S3D) and combined with conservation-based positional conditioning (*C*_*o*_) to yield joint (*C*_*o*_, *σ*_*o*_) double masks. In total, 16 residues in the REC lobe and 7 residues in the NUC lobe were fixed based on co-evolutionary coupling and mutagenesis data.

(*C*_*0*,_ *σ*_*0*_) increments were taken such that each parameter variation would result in ∼5% identity change in the generated sequences relative to the WT sequence. Following averaging the 10,000 sequence batches for each (*C*_*0*,_ *σ*_*0*_), they were manually filtered to remove redundancy, i.e. sequences having <5 mutations to the closest neighbor were removed. The resulting 43 and 44 sequences were generated independently by ESM-IF1 conditioned on the (*C*_*0*,_ *σ*_*0*_) masks for the REC and NUC lobes, respectively.

### Cloning of AI-generated TnpB variants / lobes permutation library

The generated lobes were synthesized by pairwise assembly of Oligo Pools libraries (Twist), and following addition of the WT REC and NUC lobes, cloned into custom carrier vectors via Golden Gate assembly. Finally, the resulting 1980 REC-NUC combinations variants were cloned into an expression vector with the proteins and reRNA under T7 promoters.

### Bacterial AI-generated variants library selection assay

Library transformations were performed by electroporating 0.2 ng of the TnpB expression vector library into 25 µL of *E. coli* BW25141 (λDE3) cells containing either the *ccdB* toxin plasmid (sample) or an empty control (reference). Transformants were recovered for 2 h at 37 °C with shaking and plated directly: transformations into the *ccdB*-expressing strain were plated on LB agar supplemented with chloramphenicol (25 µg/mL) and 0.2% arabinose to induce toxin expression (sample), whereas control transformations were plated on LB agar with chloramphenicol alone (reference). Colonies were scraped from sample and selection plates, and plasmid DNA was purified using the ZymoPURE II Plasmid Midiprep Kit (Zymo Research). Purified plasmids were digested with the NruI-HF restriction enzyme, and linearized fragments containing protein-coding sequences were gel-extracted and subjected to nanopore sequencing for the lobes permutation library enrichment analysis.

### Lobes permutation library enrichment analysis

We used nanopore sequencing to quantify how often each designed REC– NUC lobe combination appeared in the pooled selection plates vs reference (non-selective) plates, by mapping every nanopore read to the full set of 1980 permuted constructs (alignment templates). Briefly, raw nanopore FASTQ reads were aligned with minimap2 in ONT mode (-ax map-ont -N 5-p 0.8) against a FASTA containing all 1980 template ORFs representing the cloned AI-generated TnpB lobe permutations. For each alignment, the read was projected onto the alignment template coordinates. Then we evaluated both orientations (read vs reverse complement), retaining the orientation with the lower mismatch total. To ensure that only full-length constructs were counted, we applied two mismatch filters over functionally relevant windows (1–530 and 574–1112), each allowing ≤80 mismatches, including gaps. Alignments failing the window filter were discarded. We summed the number of passing reads *S*_*i*_ and *R*_*i*_ per template *i* for each sample and reference plates replicates, respectively. Template/expression plasmid enrichment F_*i*_ was calculated as

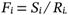

All *F*_*i*_ were normalized to the reference WT ISDra2 *F*_*i*_. Only templates/constructs *i* having *R*_*i*_ > 60 reads and *S*_*i*_ > 0 for both replicates were taken into account, and plotted in Fig. 2C.

### Mammalian cells experiments and flow cytometry

HEK293T-BFP cells (UC Berkeley Cell Culture Facility) were cultured in Dulbecco’s Modified Eagle Medium (DMEM) containing L-glutamine and 4.5 g/L D-glucose (Gibco) supplemented with 10% fetal bovine serum and 100 U/mL Penicillin-Streptomycin (Gibco). Cells were passed with Trypsin-EDTA (0.25%, Phenol Red, Gibco). Transfections were performed with 100 ng of plasmids (1:3 molar ratio of reRNA:TnpB) with TransIT LT1 Transfection Reagent (Mirus) and added to cells at ∼70% confluency. All experiments were performed in biological triplicates with the same expression plasmids used in bacterial survival assays. Genomic DNA extraction was performed using the QuickExtract solution following the manufacturer’s instructions (Lucigen). Locus specific amplification was performed with the primer sequences in table S2. Sequencing libraries were generated with Illumina adapters. All samples were quantified by qPCR using NEBQuant (NEB), pooled by equal molar amounts of each library and profiled via TapeStation (Agilent Technologies 2200).

For flow cytometry analysis, the cells were detached with 0.25% Trypsin-EDTA (Gibco), washed with PBS (Gibco), and analyzed on an Attune NxT (Thermo Fisher Scientific) for the double transfected population, positive for reRNA (mNeon+) and RNA-guided nuclease (mCherry+) population. Endogenous *BFP* levels for % Knock-out analysis were recorded. Results were interpreted using FlowJo (v10.10).

### Next-generation sequencing and data processing

Next Generation Sequencing was performed on an Illumina NextSeq 2000 instrument (P1 reagent, 600 cycles). Sequencing data processing - Read quality was assessed on raw FASTQ files with FastQC (v1.11). Bases with a quality score below 30 were trimmed from both reads using cutadapt (cutadapt -q 30). Paired-end reads were trimmed with fastp and processed using --detect_adapter_for_pe --cut_front --cut_front_window_size 4 -- cut_front_mean_quality 30 --cut_tail --cut_tail_window_size 4 --cut_tail_mean_quality 30 --qualified_quality_phred 30 --unqualified_percent_limit 30 --trim_poly_g. Trimmed paired-end reads were analyzed with CRISPResso2 (v2.3.1) (https://github.com/pinellolab/CRISPResso2)(*58*) using the reference amplicon and guide sequences (table S2). Indel frequencies were exported.

### Plant genome editing experiments

The TnpB coding sequences and overlapping reRNAs were ordered as single gene blocks with PaqCI flanking restriction sites from IDT and inserted into the plant expression T-DNA binary vector pMK525 using golden-gate cloning.

Protoplast isolation and transfection (*59*) was performed as previously described in Weiss et al (*41*). In short, protoplast cells were isolated from four week old Arabidopsis leaves. Protoplasts were then transfected with 20 µg of plasmid using polyethylene glycol (PEG)-CaCl2 solution. Following transfection, protoplast cells were incubated for 48 hours at 26 °C. During the 48-hour incubation, the protoplast cells were subjected to a 37 °C heat-shock treatment for 2 hours at 16 hours post transfection. At 48 hours post transfection, protoplasts were collected for genomic DNA extraction. Qiagen DNeasy plant mini kit (Qiagen, 69106) was used to extract genomic DNA from protoplast samples. These genomic DNA samples were then used for amplicon sequencing using the primers in table S2. Analysis of amplicon sequencing data was performed as previously described in Weiss et al(*41*) using the CrispRvariants R package (v.1.14.0).

### Protein purification

*Escherichia coli* BL21-AI cells were co-transformed with plasmids encoding 10×His–MBP–protein and T7–reRNA–HDV. A 5-mL starter culture was grown in 2× Yeast Extract Tryptone (2×YT) medium supplemented with ampicillin (100 µg//mL) and chloramphenicol (34 µg/mL) for 10 h at 37 °C with shaking at 170 rpm, then used to inoculate a 120-mL culture, which subsequently seeded 9 L of expression culture. Cells were grown at 37 °C with shaking until OD6oo ≈ 0.7, induced with 0.2% (w/v) arabinose, and incubated for 20 h at 16 °C. Cells were harvested by centrifugation and resuspended in buffer containing 20 mM Tris-HCl (pH 8.0), 250 mM NaCl, 5% (v/v) glycerol, 25 mM imidazole (pH 8.0), 1 mM TCEP, and protease inhibitor cocktail (cOmplete, EDTA-free, Roche). After sonication and ultracentrifugation, the clarified lysate was incubated with Ni-NTA resin, washed five times with buffer containing 20 mM Tris-HCl (pH 8.0), 500 mM NaCl, 25 mM imidazole, 5% (v/v) glycerol, and 1 mM TCEP, and eluted with 300 mM imidazole. The eluate was dialyzed overnight with TEV protease in buffer containing 20 mM Tris-HCl (pH 8.0), 250 mM NaCl, 25 mM imidazole, 5% (v/v) glycerol, and 1 mM TCEP. The cleaved protein was separated from the His-MBP tag using a HisTrap column (Cytiva), concentrated, and purified by ion-exchange chromatography on a HiTrap Heparin HP column (Cytiva) with a NaCl gradient from 250 mM to 1 M. The protein fraction was concentrated and further purified by size-exclusion chromatography on a Superdex 200 Increase 10/300 GL column equilibrated in cryo-EM buffer (20 mM Tris-HCl, pH 8.0, 150 mM NaCl, 2 mM MgCl2, 10 µM ZnCl2, and 1 mM DTT).

### CryoEM sample preparation and data collection

Complementary DNA oligonucleotides oSP_1437 and oSP_1438 (TS and NTS from (*32*), table S2), bearing phosphorothioate modifications, were mixed at a 45:55 molar ratio, heated to 95 °C, slowly cooled to room temperature, and purified by excision from an 8% native PAGE gel. The AI-generated nuclease was assembled with duplex DNA at a 1.12:1 (dsDNA:protein) molar ratio at 5 µM protein, incubated for 1 h at room temperature, and used immediately for vitrification. Grids (UltrAuFoil 1.2/1.3, 300-mesh gold; Electron Microscopy Sciences, #Q350AR13A) were glow-discharged at 15 mA for 25 s (PELCO easiGlow). The sample was applied to a FEI Vitrobot Mark IV system (8 °C, 100% RH), blotted for 5 s with blot force 8, and plunge-frozen. Micrographs were recorded on a Titan Krios G3 (300 kV) equipped with a Gatan K3 direct electron detector operating in CDS mode and a BioQuantum energy filter, at a nominal magnification of 105,000× in super-resolution mode (0.424 Å/pixel). Micrographs collected during seismic activity were discarded. Data was acquired using SerialEM v3.8.7 with stage- and beam-shift–based multi-shot imaging.

### Single-particle cryoEM data processing

A total of 30,156 movies were collected with a defocus range of −0.8 to −2.0 µm and processed using cryoSPARC v4.3.0. Beam-induced motion was corrected using patch motion correction, and CTF parameters were estimated with patch CTF. After micrograph curation, 15,596 images were retained for further analysis. One round of Topaz training was performed to enhance identification of the particles (*60*). From 55 representative micrographs, 39,091 particles were initially picked with the blob picker and subjected to 2D classification, and 2,487 high-quality particles from the best classes were used to train a Topaz model. This model was then applied to all curated micrographs, yielding 3,216,718 particle picks (binned twofold) that were classified in 2D. From the best classes, 1,496,915 particles were selected for *ab initio* reconstruction and heterogeneous refinement into two classes. The 955,983 particles forming the best-resolved class were refined using non-uniform refinement. These particles were subsequently re-extracted and subjected to an additional round of *ab initio* reconstruction, heterogeneous refinement, and final non-uniform refinement, resulting in two distinct maps corresponding to the TAM-bound and R-loop conformations. Each map, reconstructed from 389,833 and 566,150 particles respectively, achieved a resolution of 2.8 Å and was further sharpened using DeepEMhancer prior to model building. The corresponding statistics are summarized in Table S1.

### Single-particle cryoEM model refinement

The initial model of the AI-generated protein v7 was obtained with AlphaFold2, whereas 8BF8 and 8EXA from (*34*) were used for the nucleic acids. Following morphing in Coot v0.9.4.1, the models were refined using the real-space refinement and rigid body fit tools. RNA coordinates were optimized with the ERRASER tool (*61*). Finally, the model was subject to a round of real_space_refine tool in Phenix v1.21.1-5419. Cryo-EM maps and model coordinates were deposited to the EMDB (codes EMD-73644, EMD-73645) and PDB (codes 9YYG, 9YYH). The structures’ figures were generated in UCSF ChimeraX v1.5.

